# Lipidome profiles of postnatal day 2 vaginal swabs reflect fat composition of gilt’s postnatal diet

**DOI:** 10.1101/593392

**Authors:** KaLynn Harlow, Christina R. Ferreira, Tiago J.P. Sobreira, Theresa Casey, Kara Stewart

## Abstract

We hypothesized that postnatal development of the vagina is impacted by early nutritional environment. Our objective was to determine if lipid profiles of vaginal swabs were different between gilts suckled by sow or fed milk replacer the first 48 h postpartum, with and without a lard-based fat supplement. Gilts (>1.3 kg) were selected at birth across 8 litters and assigned to treatments: colostrum suckled (S, n=8); S plus fat supplement (SF, n=5); bottle-fed milk replacer (B, n=8); or B plus fat supplement (BF, n=7). At 48 h postnatal, vaginal swabs were taken with a cytology brush, immersed in ultrapure water to burst cells, and lipids extracted for analysis using multiple reaction monitoring (MRM)-profiling. Lipids extracted from serum collected at 48 h from gilts and milk collected from sows at 24 h were also analyzed with MRM-profiling. Receiver operating characteristic curve analysis found 18 lipids highly distinguished [area-under-the-curve (AUC) > 0.9] between S and B gilts, including phosphatidylethanolamine with 34 carbon and four unsaturations in the fatty acyl residues [PE(34:4)]. Twelve lipids from vaginal swabs highly correlated (*r* > 0.6; *p* < 0.01) with nutrition source. Lipids more abundant in milk replacer drove association. For example, mean intensity of PE (34:4) was 149-fold higher in milk replacer than colostrum, with 1.6- and 2.12-fold higher levels in serum and vaginal swab samples (*p* < 0.001), respectively, of B versus S gilts. Findings support that vaginal swabs can be used to noninvasively study effects of perinatal nutrition on tissue composition.

**Summary sentence:** Vaginal swab lipidome profiles at 48 h reflect the fat composition of neonatal diet during first two days postnatal.

## Introduction

Early nutritional environment affects long term health and fertility. In swine, colostrum ingestion is essential for postnatal piglet survival, growth, and development because it provides immunity, nutrients, energy, and bioactive factors [1, 2]. The window of opportunity for milk-borne bioactive factors to influence neonate development is limited, and primarily occurs prior to closure of tight junctions between cells lining the piglet’s gut. Closure of the gut occurs by 48 h postnatal [3]. During the first 48 h postnatal, piglets ingest up to 30% of their body weight in milk [4]. This time-period is a critical developmental period for the gilt reproductive system, including the formation of uterine glands, otherwise known as adenogenesis [5, 6]. Colostrum ingestion significantly affected the developmental trajectory of uterine tissue [7–10]. Replacement gilts with less colostrum consumption than littermates as indicated by blood immunocrit values had reduced litter sizes relative to other sows [11, 12]. Colostrum-deprivation also resulted in significantly different patterns of uterine gene and protein expression [13].

The link between early nutritional environment, uterine development, and subsequent reproductive potential led to the hypothesis that early nutritional environment affects reproductive tract development and subsequently predicts long-term reproductive performance of gilts. However, in order to evaluate uterine development, the animal must be euthanized. We previously proposed that since the vagina is embryologically related to the uterus [14], its postnatal developmental trajectory may also be responsive to early nutritional environment. Moreover, we proposed that using vaginal swabs to non-invasively sample lower reproductive tract may serve as a means to evaluate differences in nutritional exposures on gilt development. Using a biomarker-discovery technique known as multiple reaction monitoring (MRM) profiling, we found that lipid profiles of vaginal swabs taken on postnatal day 14 differed between gilts that were fed milk replacer during the first 48 h postpartum before return to litter versus gilts that suckled sow’s milk continuously from birth [15].

While our previous studies supported the potential of using biological material obtained from vaginal swabs to distinguish between gilts exposed to different nutritional environments the first 2 days postnatal [16], the lapse of time from colostrum exposure and relatively small sample size limited interpretation. In this study, we further investigated the efficacy of using MRM-profiling of vaginal lipids to differentiate PND 2 vaginal swabs between gilts suckled by sow or fed milk replacer. Secondly, we tested the effect of a lard based supplement on vaginal lipid profiles of gilts [17–20].

## Materials and Methods

### Animals and study design

Prior to beginning studies involving animals the protocol was reviewed and approved by Purdue University’s Institutional Animal Care and Use Committee (Protocol #1605001416). Standard farrowing protocols for the Purdue University Animal Sciences Research and Education Swine facility were followed. All lipidomics sample preparation and analysis was completed at the Proteomics and Metabolomics Core Facilities in the Bindley Bioscience Center at Purdue University.

Three to four gilts were selected per litter from eight different sows which were monitored during parturition (Supplemental Figure S1). Immediately after delivery, all gilts were towel-dried, weighed, and placed in a holding cart until at least three gilts above 1.3 kg were delivered. Within litter, each gilt was randomly assigned to one of four treatment groups and ear tagged for identification. The four treatment groups were: suckled (S; n=8); suckled plus fat-supplement (SF; n=5); bottle-fed with milk-replacer (B; n=8), and; bottle-fed milk replacer plus fat-supplement (BF; n=7). Body weights were recorded at birth and 48 h. All gilts were administered a 2 ml dose of Camas experimental antibody product (Camas Incorporated; Le Center, MN) using Pump It™ Automatic Delivery System (Genesis Industries, Inc.) at birth, and at 3 h and 9 h after the first dose.

Gilts in B and BF treatment groups were taken to a separate temperature-controlled nursery, where they were placed in holding cages in groups of 2-3 piglets. Birthright™ milk replacer (Ralco Nutrition, Inc.) was mixed using the recommended concentration for orphaned piglets, which was 1:2 replacer to tap water. Reconstituted milk replacer was stored in a refrigerator and warmed to room temperature prior to feedings. Piglets were fed using Evenflo Classic slow-flow 0-3-month infant bottles (Evenflo Feeding, Inc.) approximately every 2 h, with a minimum of 5 ml fed per time point. In the event piglets did not consume at least 30 ml after two consecutive feedings, 15 ml of milk replacer was oral gavage-fed using a 30 ml syringe and Kendall™ 4.7 mm x 41 cm feeding tube (Medtronic, Minneapolis, MN). The fat supplement was administered at the back of the throat in 3 ml room-temperature doses using a 5 ml syringe at 6 h, 12 h, 24 h, and 36 h after birth. Swallowing was observed to ensure the supplement was delivered.

Blood samples were collected from gilts at 48 h postnatal. A 22 g x 1.5-inch needle and a 3 ml serum Vacutainer^®^ (BD Life Sciences) tube was used to collect blood by jugular venipuncture. Blood samples were refrigerated and allowed to clot overnight. Serum was collected following centrifugation (Horizon Model 614B Centrifuge; Fisher Scientific) for 25 minutes at 2,500 g and stored at −20°C until analysis.

Gilts were euthanized approximately 48 h after birth using CO_2_ inhalation. Following euthanasia, skin of the abdominal and genital regions was cleaned thoroughly using 70% ethanol, and vaginal swabs were taken using a cytology brush (Puritan 2196 Removable Stiff Bristle Tip Brush; QuickMedical; Issaquah, WA) by inserting the tip of the brush into the vulva angled dorsally at 45°. Once inserted to the base of the bristles, the brush was rotated 360° against the vaginal surface. Two consecutive swabs were collected from each animal, and swabs were placed in separate 15 ml sterile polypropylene conical tubes (Falcon™, Fisher Scientific, San Jose, California) and immediately placed on dry ice for transport. Samples were stored in a −80°C freezer until lipid extraction and analysis.

### Fat supplement preparation

The fat supplement was created by emulsifying pork lard (Berkshire pork lard, EPIC; Austin, TX). Pork lard was used as the fat source because the fatty acid composition of lard is similar to sow colostrum [21]; Tween 80 was used as an emulsifying agent. A 30% volume/volume solution of lard/2% aqueous solution of Tween 80 was prepared using a homogenizer to microscopically distribute the lipid aggregates with low heat. The emulsion was stored at 4°C.

### Milk collection

Sows were milked during farrowing, and at 24 h after delivery of first piglet. For milk collection piglets were removed from the sow for approximately an hour and then 1 ml oxytocin (VetOne; Boise, ID; 20 USP/ml) was administered IM using a 20g x 1.5-inch needle into the vulva to stimulate milk letdown. Colostrum samples were collected manually from all teats and combined to create a uniform sample. Samples were stored until further analysis at −20°C.

### Lipidomics Analysis

#### Lipid extraction

The Bligh & Dyer [22] lipid extraction technique was slightly modified to extract lipids from the vaginal swab samples. A volume of 500 μl distilled water was added to the conical tube containing the swab brushes and vortexed to remove biological material from the brush. The brushes were removed, the sample homogenate was transferred to a new tube, and phase separation was performed by mixing with chloroform/methanol/distilled water (1:2:0.8).

Samples were centrifuged, the organic phase (bottom phase) was separated from aqueous phase, divided into four aliquots, and dried in a centrifugal evaporator (Savant SpeedVac AES2010, ThermoFisher Scientific, San Jose, CA). Dried lipid extracts were stored at −20°C until mass spectrometry analysis.

The Bligh & Dyer [22] lipid extraction technique was also used to extract lipids from 48 h serum samples from suckled and bottle-fed piglets; colostrum samples taken from all eight sows (during parturition, 6 h after first piglet born, 12 h after, and 24 h after); and milk replacer used for bottle-feeding. The procedure for these samples began at the phase separation step (i.e. no water was added to the samples).

#### Multiple reaction monitoring (MRM)-profiling

For the discovery phase of MRM-profiling, vaginal lipid extracts were pooled by treatment to determine the functional groups present in samples. Pooled samples were prepared by combining equal volumes of dried lipid extracts diluted in 200 μl of the flow injection solvent [acetonitrile/methanol/ammonium acetate 300 mM at 3:6.65:0.35 (*v*/v)]. Pooled sample injections (8 μl) were delivered to the micro-autosampler (G1377A) in a QQQ6410 triple quadrupole mass spectrometer (Agilent Technologies, San Jose, CA) equipped with an ESI ion source. The solvent pumped between injections (CapPump G1376A, Agilent Technologies, San Jose, CA) was acetonitrile with 1% formic acid at 10μL/min. Between sample injections, methanol/chloroform was injected to remove any remaining lipids before the next sample injection. Samples were examined for fragmentation features of specified chemical classes using Precursor (Prec) and neutral loss (NL) scan modes as previously reported [15]. Scans were used to profile chemical classes, which included: phosphatidylethanolamine (PE), acylcarnitines (AC), cholesteryl esters (CE), glycosylinositolphosphoceramides (GIPC), phosphatidic acids (PA), phosphatidylcholines (PC), alkenyl-acyl PCs (ePC), sphingomyelin (SM), phosphatidylinositols (PI), phosphatidylglycerols (PG), phosphatidylserines (PS), fatty acids (FA), ceramides, sugar lipids, and glycerolipids (see Supplemental Table S1 for classes scanned). Initial data processing of the profiles obtained was carried out by converting each set of MRM-profiling method data into mzML format using MSConvert20. Next, the signal intensity for the ions present in NL and Prec mass spectra was obtained using an in-house script. Ions with values of counts >1000 were selected as parent ions and the product ion or neutral loss information was used for selecting ion pairs (1486) for the screening phase.

For Screening Phase I, individual sample extracts were analyzed for the MRMs identified in the discovery phase (1486), as well as triacylglycerol (TAG) MRMs (Supplemental Table S2). TAG have no polar head or diagnostic functional group fragment, precluding them from the discovery step of analysis. During screening phase, TAG were profiled using parent ions and a product ion related to the presence of specific fatty acyl residues (16:0, 16:1, 18:0, 18:1, 18:2, and 20:4). MRMs identified in the discovery step and TAG MRMs were divided into 8 groups, or 8 screening *methods*. The methods were organized by factors such as lipid class to optimize the efficiency of the machine and to limit the number of precursor ion/product ion pairs screened per sample injection. Thus, for each of the 8 methods, intensities of approximately 250 MRMs were measured per 8μl sample injection. Since cellular content in each sample may vary when using swabs, prior to screening lipid extract amount was recorded across samples by scanning for the precursor ion of *m/z* 184, which profiles the most abundant lipids in cells, namely PC and SM lipids. In this way, intensities for PC lipids were measured, and samples were diluted using the same flow injection solvent to achieve a similar concentration for comparative relative analysis.

Data from Screening Phase I were uploaded into MetaboAnalyst 4.0 (www.metaboanalyst.ca) to identify MRMs that best discriminated between suckled and bottle-fed gilts. A second set of MRMs were selected based on discrimination between gilts with and without fat supplementation. There were 146 MRMs identified that best discriminated between suckled and bottle-fed vaginal lipids (S and SF piglets versus B and BF piglets, respectively), and 197 MRMs that discriminated not-supplemented and fat supplemented vaginal lipids (S and B piglets versus SF and BF piglets, respectively). The 146 MRMs (Supplemental Table S3) and 197 MRMs (Supplemental Table S4) were organized into two distinct methods (Method 1 and Method 2, respectively) for Screening Phase II analysis.

Method 1 and Method 2 were used to evaluate the vaginal lipids from all piglets in Screening Phase II. The resulting ion pair intensities were normalized to reflect percent intensity for subsequent comparative analysis. These data were used to verify the effectiveness of each method to differentiate between piglets that were: A.) suckled or bottle-fed, or; B.) fat supplemented or not. Data were uploaded in MetaboAnalyst 4.0 by method for statistical analysis to evaluate the potential effectiveness of selected MRMs as biomarkers for distinguishing treatment groups.

#### MRM-profiling data analysis in MetaboAnalyst

Ion intensities of MRMs from the screening phase were used to calculate the relative ion intensity of ion pairs in each sample and these data was uploaded into MetaboAnalyst 4.0 [23] and normalized using autoscaling. The Biomarker Analysis tool was used to identify MRMs that discriminated between suckled versus bottle-fed gilts and fat supplemented versus non-fat supplemented gilts. Biomarker analysis tools applied included principal component analysis, partial least square-discriminant analysis, and classical univariate receiver operating characteristic (ROC) curve analysis. Using data from Screening Phase I, a cut-off area-under-the-curve (AUC) value of 0.70 and 0.80 was used to select MRMs to discriminate between vaginal lipids of piglets suckled versus bottle-fed (Method 1), and those with and without fat supplementation (Method 2).

#### Lipid identification

Precursor ion/product ion pairs were tentatively assigned identities using attributions in Metlin (https://metlin.scripps.edu) and Lipid Maps (http://www.lipidmaps.org/). Potential identities were assigned using the associated functional group and biological information. Metlin MS/MS Spectrum Search was used with the following settings: precursor ion mass from the mass spectrometer reading for *m/z* value, maximum tolerance at 100, collision energy at 10 eV or 20 eV, MS/MS tolerance at 0.5, peaks included the product ion mass *m/z* with an intensity of 100, positive mode, and M+H was selected for the adduct. If the suspected ion pair was a TAG, M+NH_4_ was also selected as an adduct to account for ammonium acetate.

Phospholipids were assigned identity by their class (PS, PI, PE, or PC), the number of carbon atoms in the esterified fatty acid, and the number of carbon-carbon double bonds present in the molecule, eg. PE(34:4). TAGs were similarly assigned tentative attributions. First, the number of carbon atoms in each fatty acyl chain was summed, and the number of carbon-carbon double bonds was assigned. Since searches in Lipid Maps can only be performed using neutral masses, and ammonium acetate was present in the injection solvent, TAGs were detected as ammonium adducts. For this reason, 18 mass units (NH_4_) was subtracted from the parent mass observed in the MS analysis to obtain the neutral mass for each TAG. Other lipids were searched by accounting for a single mass unit loss to account for a hydrogen ion only. Results were narrowed using the associated functional group and biological information, and attributions were assigned where possible.

#### Swab, serum, and colostrum correlation analysis

Method 1 was also used to analyze serum lipids collected from the S and B animals, milk replacer, and colostrum at 0 h, 6 h, 12 h, and 24 h post-parturition. These resulting intensities were converted to percent intensity for comparative analysis. Pearson’s correlation coefficient analysis of percent intensities between milk and serum samples, milk and vaginal swab samples, and serum and swab samples was performed for each of the 38 most discriminating MRMs from Method 1 using the CORR procedure of SAS 9.4 (Cary, NC). Variables compared were percent intensities of: 48 h swab samples and 48 h serum samples; 48 h swab samples and respective source of nutrition at 24 h; and 48 h serum samples and respective source of nutrition at 24 h. Six MRMs overlapped as highly correlating between source of nutrition and swab lipids, and source of nutrition and serum lipids. Percent ion intensities were compared between milk replacer and colostrum for those six MRMs. Statistics were not performed on colostrum versus milk replacer since ion intensity results were from only one sample of milk replacer. In addition, MIXED procedure of SAS 9.4 (Cary, NC) was used to analyze the statistical difference in ion intensities of the six most highly correlated MRMs between suckled versus bottle-fed gilt serum, as well as suckled versus bottle-fed gilt swab samples to compare.

## Results

### Effect of postnatal diet on vaginal lipid profiles

The discovery phase identified a total of 1486 unique MRMs in pooled isolates from vaginal swabs. Glycerolipids with saturated bonds encompassed 37% of identified lipids (Figure 1A; Supplemental Table S2). Glycerolipids with unsaturated bonds were next in abundance with 25% of identified lipids, and acylcarnitines were third most abundant (15.81%). Intensities of the 1486 MRMs identified in the discovery phase, plus approximately 400 triacylglycerols (TAGs), were measured in individual samples in Screening Phase I (Supplemental Table S2). Screening Phase I analysis found 146 MRMs (Supplemental Table S3) that best discriminated between suckled and bottle-fed vaginal lipids, and 197 MRMs (Supplemental Table S4) that best discriminated between vaginal lipids of gilts with and without fat supplementation.

**Figure 1.**
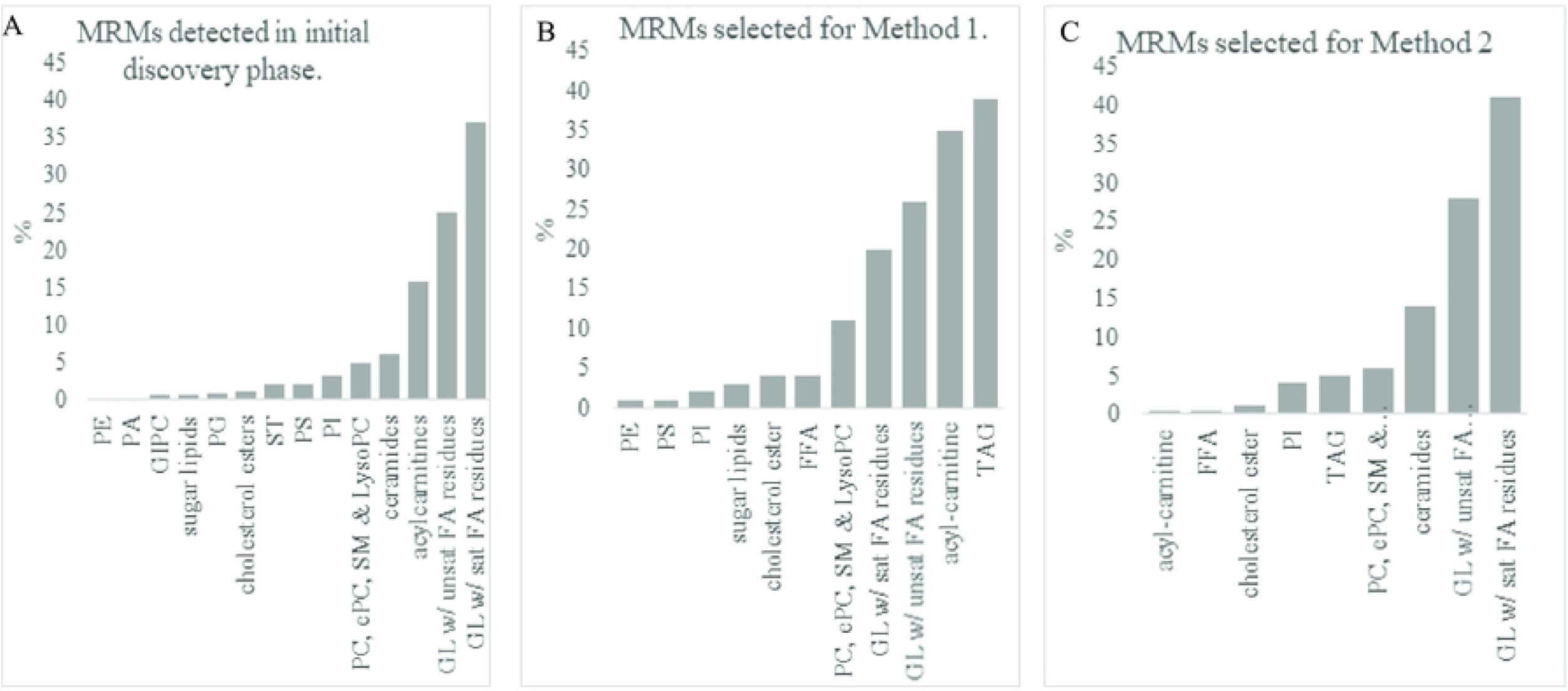
Lipid class distribution of MRMs: (A) detected in initial discovery phase; (B) Selected for method 1; and (C) Selected for method 2.

For Screening Phase II, the 146 MRMs that distinguished between suckled and bottle-fed treatments were organized into Method 1 analysis, which consisted primarily of TAGs, acylcarnitines, and glycerolipids (Figure 1B). Tentative attributions were assigned to 91/146 lipids in Method 1 (Supplemental Table S5). Principal component and hierarchical cluster analysis showed distinct clusters by treatment groups of suckled versus bottle-fed milk replacer (Figure 2A and 2B). ROC curve analysis identified 18 MRMs with AUC values greater than 0.90. Of the 18, 15 were assigned tentative attributions, which included hydroxy octadecanoyl carnitine, two TAG(46:1), TAG(48:2), TAG(48:1), PG(36:2)/PI(30:4), PC(36:7), TAG(44:1), PG(34:6), PE(34:4), PE(36:5), PE(38:5), two TAG(44:0), and TAG(48:3). The proportion of TAGs identified in this group (44%) was greater than proportion of TAGs present in the entire list of Method 1 ion pairs (27%). The 20 most discriminating lipids recovered from the vaginal swabs showed a greater relative abundance in most bottle-fed versus suckled gilts (Figure 2B).

**Figure 2.**
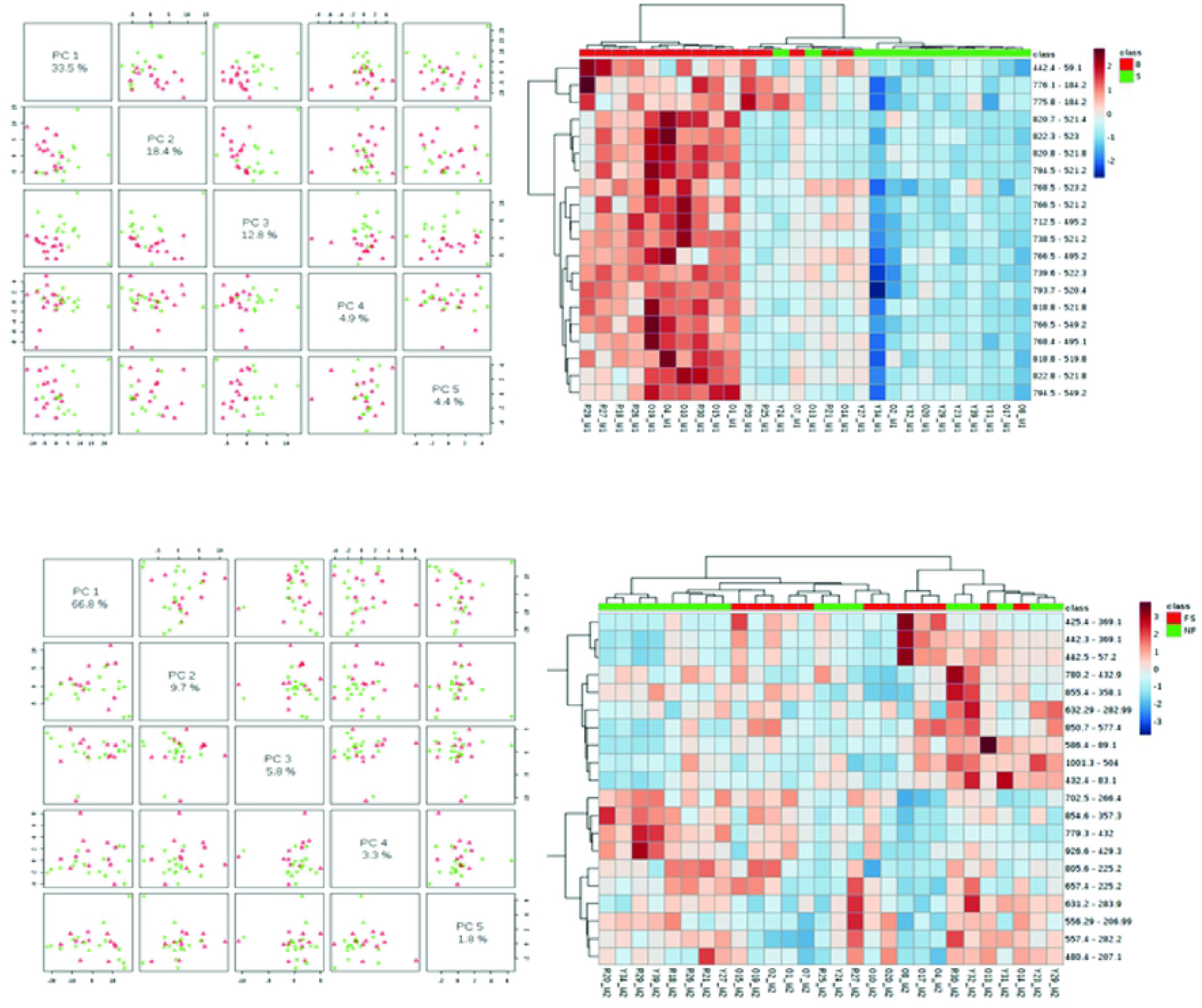
(A) Principal component analysis and (B) heat map of lipids selected for Method 1; within the hierarchical analysis, red signifies bottle-fed animals (B and BF) and green signifies suckled animals (S and SF). (C) Principal component analysis and (D) heat map of lipids selected for Method 2; within the hierarchical analysis, red signifies fat supplemented (SF and BF) and green signifies not supplemented (S and B) animals. In the heatmap, blue indicates lower ion intensities and red indicates higher ion intensities. MRMs are on the right and animal ID is on the bottom.

Hierarchical cluster and principal component analysis of the effect of fat supplementation on lipidome profiles showed several groupings, with some distinction between plus and minus fat supplemented gilts (Figure 2C and 2D). The 197 MRMs that most distinguished between fat supplemented piglets and non-supplemented piglets and selected for Method 2 Screening Phase II analysis were primarily glycerolipids (Figure 1C). ROC curve analysis identified 15 MRMs with AUC values ranging from 0.65 to 0.78. Mean ion intensity of these MRMs was compared between treatments, and overall treatment effects (*p* < 0.05) were found for a cholesteryl ester and a ceramide, with numerical differences between treatments in two glycerol lipids with EPA residues (Figure 3).

**Figure 3.**
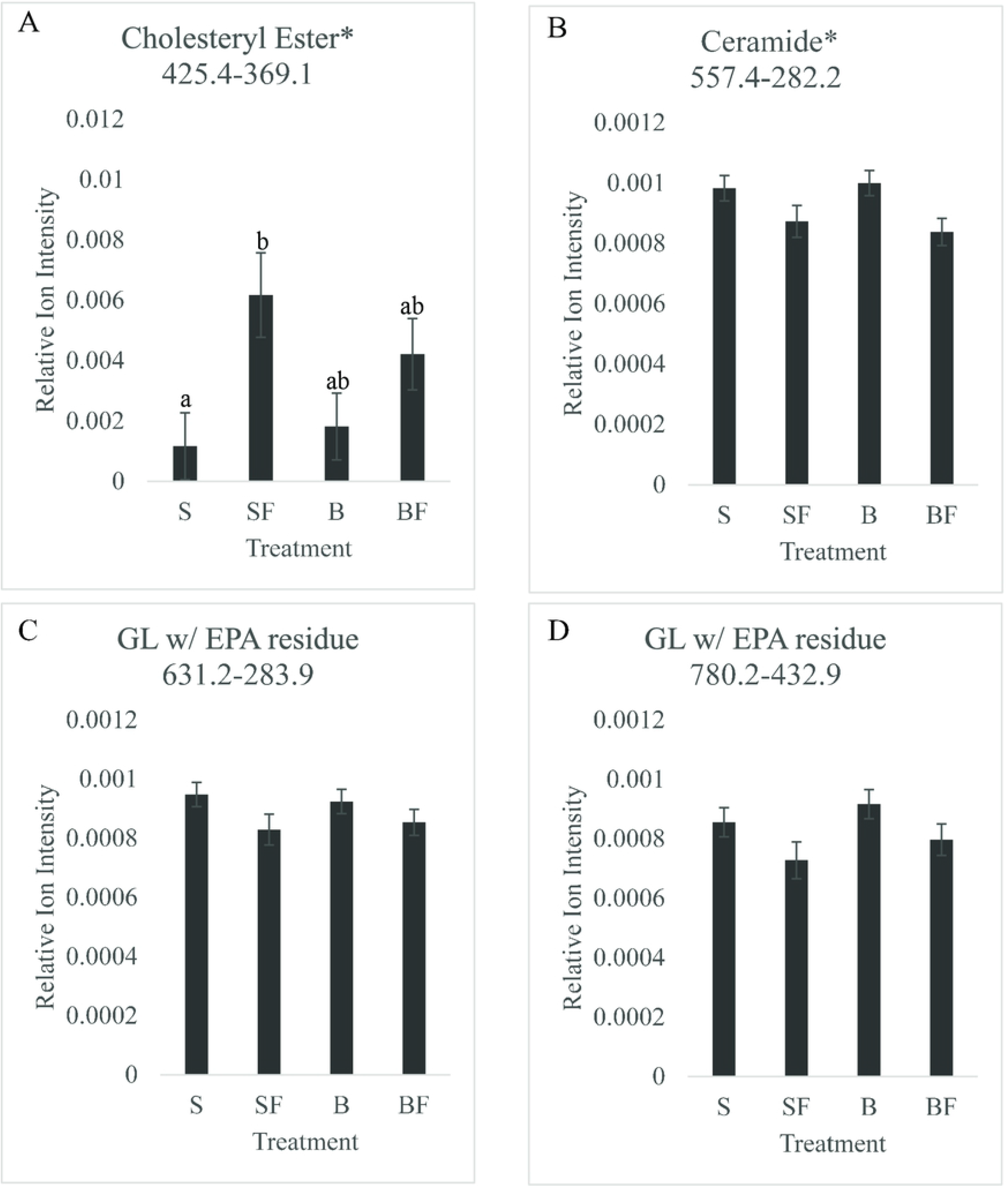
Mean ion intensity of precursor-product ion MRMs of (A) cholesteryl ester 425.4->369; (B) ceramide 557.4->282.2; (C) glycerolipid containing eicosapentaenoic acid residue 631.2->283.9; and (D) glycerolipid containing eicosapentaenoic acid residue 780.2->432.9 measure in lipid extracts of vaginal swabs taken from bottle fed (B), bottle fed and fat supplemented (BF), suckled by sow (S) and suckled by sow plus fat supplemented (SF) for 48 h postnatal. Differing letters indicate statistical difference at *p*<0.05; and (*) indicates an overall fat supplement treatment effect.

### Correlation of lipids isolated from Swabs, Blood Serum, and Colostrum-Milk Replacer

To investigate whether there was a relationship between piglet nutrition and either serum or vaginal lipidome profiles, lipids from piglet serum, sow colostrum, and milk replacer were analyzed using Method 1 MRMs. Linear regression analysis was used to determine if there was a relationship between intensities of the top 38 MRMs (AUC > 0.8) that discriminated most between vaginal swab samples of S and B groups with intensity in piglet serum and nutrition source (sow colostrum or milk replacer; Table 1). The correlation coefficient between twelve MRMs in swab and milk samples was highly significant (r > 0.60 and *P* < 0.01), and included: three PE lipids; a PG lipid; a PC lipid; five TAGs; stearoylcarnitine, hexadecanedioic acid mono-L-carnitine ester; and PG (36:2)/PI(30:4). Analysis of relationship of lipid intensities by nutrition source and serum found seven MRMs were significant including three PE lipids, one PG lipid, and three TAGs. One MRM, related to TAG (48:3), had a negative correlation between nutrition source and serum (r = −0.62 and *P* = 0.01).

Six MRMs overlapped amongst the strongest correlations between source of nutrition and serum and between source of nutrition and swab [tentative lipid attributions: PE (34:4), PE(36:5), PG(34:6), TAG(46:1), PE(38:5), and TAG(44:0)]. Levels of all six lipids were 159-fold to 2.41-fold (Figure 4 and Supplemental Figure S2) higher in milk replacer than colostrum, which correlated to significantly higher levels across all serum and swab samples of botte-fed versus suckled animals (Figure 4).

**Figure 4:**
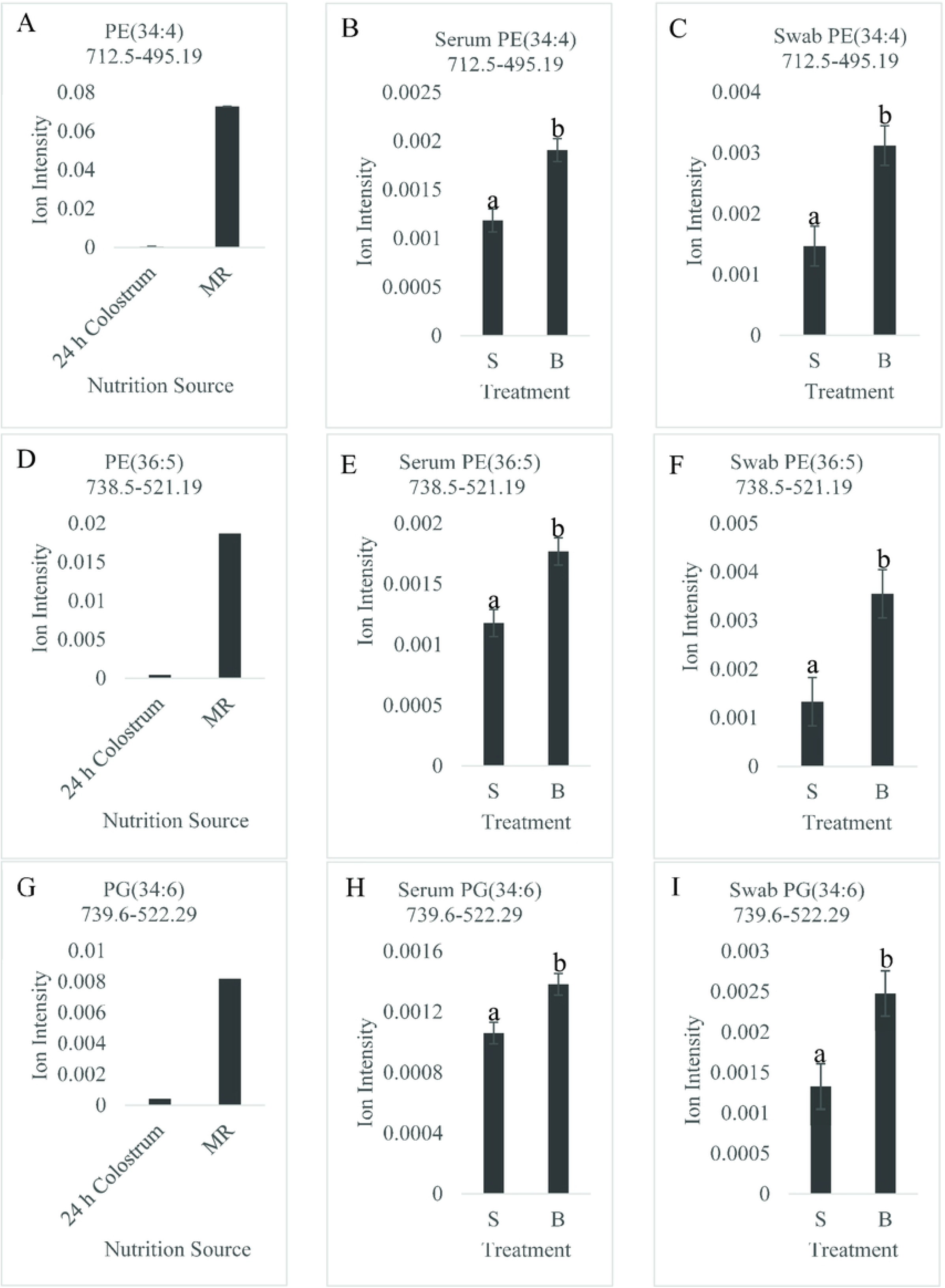
Comparison of relative mean intensity of PE (34:4), 712.5->495.19 in (A) nutrition source (24 h colostrum sample versus milk replacer-MR); (B) serum of suckled-S versus bottle fed milk replacer-B; and (C) vaginal swab samples in S versus B gilts. Relative mean intensity of PE (36:5), 738.5->521.19 in (D) nutrition source (24 h colostrum sample versus milk replacer-MR); (E) serum of suckled-S versus bottle fed milk replacer-B; and (F) vaginal swab samples in S versus B gilts. Relative mean intensity of PG (34:6), 739.6->522.29 in (G) nutrition source (24 h colostrum sample versus milk replacer-MR); (H) serum of suckled-S versus bottle fed milk replacer-B; and (I) vaginal swab samples in S versus B gilts. Differing letters indicate statistical difference at *p*<0.005.

## Discussion

Screening phase II Method 1 analysis aimed at identifying lipids that discriminated between gilts which suckled sow colostrum versus gilts fed milk replacer the first 48 h postnatal. ROC curve analysis found 18 lipids with AUC ≥ 0.9. The AUC value is a measure of the usefulness of a test, in terms of sensitivity and specificity [24]. Biomarkers with AUC values from 0.9-1.0 are excellent, 0.8-0.9 are good, 0.7-0.8 are fair, and 0.6-0.7 are poor [25]. The 18 lipids are thus excellent candidate markers to detect differences between suckled gilts compared to those that consumed only milk replacer. The most discriminatory lipids identified in Method 1 exhibited higher intensities in bottle-fed animals when compared to suckled gilts. We found a positive correlation of ion intensities of MRMs between nutrition source and swabs of suckled versus bottle-fed gilts, and nutrition source and serum of suckled versus bottle-fed gilts. Comparative analysis of mean ion intensities of the most highly correlated lipids found much higher intensities of lipid signals in milk replacer than in 24 h colostrum samples, and correspondingly higher intensities in serum and vaginal swabs of bottle-fed animals relative to suckled gilts.

The high correlation of MRM intensities between nutrition source and swab were primarily driven by lipids that were significantly more abundant in milk replacer than colostrum samples. These findings demonstrate that lipid content in milk replacer formulas changes the composition of neonate tissues, and component fats may become part of the tissues. Lipids are bioactive molecules, and play a central role in metabolism and influence and modulate cell function. For example, lipids from the diet function in cytokine and (steroid) hormone synthesis, cell differentiation and growth, cell membrane structure, myelination, signal transmission, and are substrates used for prostaglandin synthesis. Fatty acids and their metabolites, such as eicosanoids, have a major signaling function and regulate gene transcription by serving as transcriptional co-activators, with polyunsaturated fatty acids activating the transcription factor peroxisome proliferator–activated receptors [26, 27]. Thus, fatty acid composition of diet has the potential to affect significant biological response in the neonate, which in turn can affect the course of neonatal development and health. In humans, there is evidence that the quality of dietary lipids provided to infants has a marked impact on health outcomes [28], with our study supporting that dietary lipids became components of tissues. Although infant formula has been amended some, currently available products continue to be markedly different from breast milk in their lipid composition; these differences might be of importance for infant health, development and long-term fertility [28]. Correlation of lipids in nutrition source (milk replacer and colostrum) with serum and swab, suggest some lipids may be directly absorbed without modification in the gastrointestinal tract, potentially prior to intestinal barrier closure, which is within 36-48 h after birth [3, 29–31]. Thus this period of time may be viewed as either a window of opportunity or vulnerability for influencing neonate’s developmental program.

Functional groups of Screening Phase II Method 2 analysis were primarily glycerol lipids with both saturated and unsaturated FA residues. Of the most discriminatory lipids, four MRMs had AUC values between 0.7 and 0.8, thus achieving a biomarker utility rating of fair [25]. Two of these were glycerol lipids with eicosapentaenoic acid (EPA) residues, one was a ceramide, and the final was a cholesteryl ester. Mean intensity of the cholesteryl ester was significantly higher in suckled plus fat supplemented gilts versus non-supplemented gilts. Unfortunately, MRM-profiling was not completed with the fat supplement. Thus we can only speculate that the higher cholesteryl ester content in swabs was a response to the higher fat diet. Two glycerol lipids with EPA residues and ceramide were numerically lower in vaginal swabs of fat supplemented gilts. EPA is either absorbed from the diet or synthesized from the essential fatty acids linoleic and alpha linoleic acid. A potential explanation of these findings is that direct supplementation of piglets with animal fats lowered the bioavailability of EPA (or linoleic and alpha linoleic acids). In the mature gut, triglycerides and phospholipids are hydrolyzed by pancreatic lipase, but the ethyl ester of EPA requires additional digestion with bile salt–dependent lipase [32]. Digestion and absorption of EPA is thus considerably lower than triglycerides and phospholipids, and dependent on high-fat content in meals, which stimulates bile-salt dependent lipase and enhanced absorption [32]. In porcine and human neonates the levels of pancreatic lipase and bile acids are much lower than at maturity [33]. Thus, despite the higher lipid content of fat supplement treated gilts, the low availability of bile acids levels may have limited EPA absorption. Further research into how lipids ingested by early postnatal gilts are metabolized and used in the reproductive tract is needed.

## Conclusion

Vaginal swab lipidomes were significantly affected by composition of early postnatal diet. Multiple lipid molecules were differentially abundant between gilts exposed to colostrum versus gilts fed milk replacer and gilts fed with and without fat supplement the first 48 h of life. Several lipids collected from vaginal swabs were highly correlated with the nutrition source a piglet ingested. Lipids more abundant in milk replacer appeared to be driving the association in the current study. These findings demonstrated that lipid composition of neonatal diet affects tissue composition and thus may affect development, function and overall health and fertility.

Moreover, while data support the efficacy of using MRM-lipid profiling of vaginal swabs to detect differences in early postnatal nutritional exposures, findings also highlight a need to modify our approach to determine if biomarkers of colostrum intake can be specifically identified in vaginal swab samples.

**Table 5.**
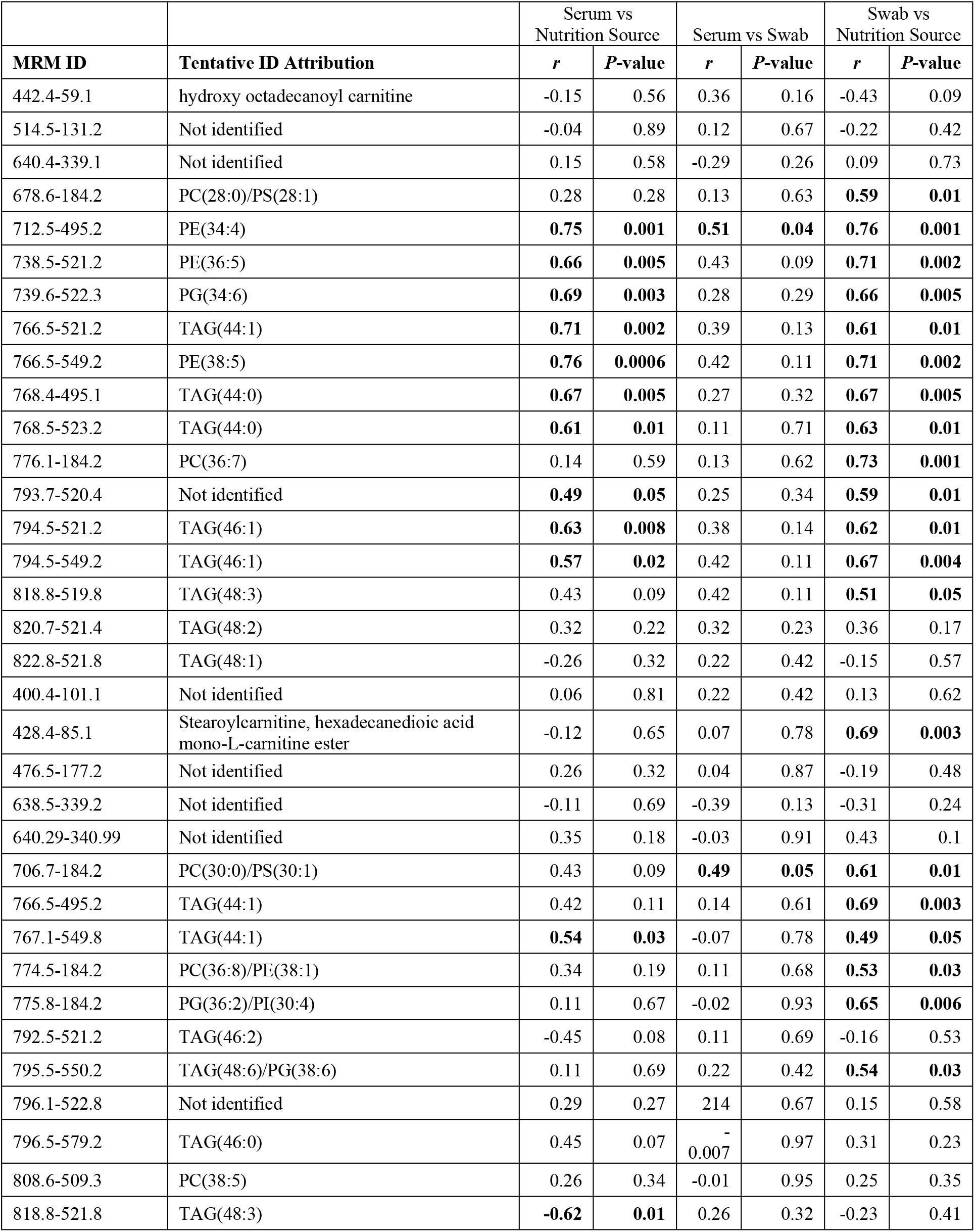
Pearson correlation coefficients of 38 MRMs with highest AUC values. Bold values indicate either r > |0.60| or P-value < 0.01.

## Abbreviations

(Prec): Precursor
(NL): neutral loss
(PE): scan modes. phosphatidylethanolamine
(AC): acylcarnitines
(CE): cholesteryl esters
(GIPC): glycosylinositolphosphoceramides
(PA): phosphatidic acids
(PC): phosphatidylcholines
(ePC): alkenyl-acyl PCs
(SM): sphingomyelin
(PI): phosphatidylinositols
(PG): phosphatidylglycerols
(PS): phosphatidylserines
(FA): fatty acids
(TAG): triacylglycerol
(B): bottle fed milk replacer
(S): suckled colostrum
(BF): bottle fed milk replacer and supplemented with fat
(SF): suckled and supplemented with fat
(ROC): receiver operating characteristic
(AUC): area-under-the-curve
(MRM): multiple reaction monitoring
(PND): postnatal day

## Supplemental Data Legends

**Supplemental Figure S1.** Study design and work flow.

**Supplemental Figure S2.** Comparison of relative mean intensity of TAG(46:1) in (A) nutrition source (24 h colostrum sample versus milk replacer-MR); (B) serum of suckled-S versus bottle fed milk replacer-B; and (C) vaginal swab samples in S versus B gilts. Relative mean intensity of TAG(44:0) in (E) nutrition source (24 h colostrum sample versus milk replacer-MR); (F) serum of suckled-S versus bottle fed milk replacer-B; and (G) vaginal swab samples in S versus B gilts. Relative mean intensity of PE (38:5) in (H) nutrition source (24 h colostrum sample versus milk replacer-MR); (I) serum of suckled-S versus bottle fed milk replacer-B; and (J) vaginal swab samples in S versus B gilts. Differing letters indicate statistical difference at *p*<0.005.

**Supplemental Table S1:** Neutral Loss and Precursor Scan Modes Used for Discovery of MRMs

**Supplemental Table S2:** MRMs Selected After Discovery Step For 8 Methods

**Supplemental Table S3.** Selected MRMs for Method 1 (Bottle-fed versus Suckled)

**Supplemental Table S4.** Selected MRMs for Method 2 (Fat supplemented versus Not fat supplement)

**Supplemental Table S5.** Tentative Attributions for Method 1

## REFERENCES

1. Peaker M. The mammary gland in mammalian evolution: a brief commentary on some of the concepts. Journal of mammary gland biology and neoplasia 2002; 7:347–353.

2. Dividich JL, Rooke JA, Herpin P. Nutritional and immunological importance of colostrum for the new-born pig. J. Agric. Sci. 2005; 143:469–485.

3. Bartol FF, Wiley AA, Bagnell CA. Relaxin and Maternal Lactocrine Programming of Neonatal Uterine Development. Annals of the New York Academy of Sciences 2009; 1160:158–163.

4. Bagnell CA, Yan W, Wiley AA, Bartol FF. Effects of Relaxin on Neonatal Porcine Uterine Growth and Development. Annals of the New York Academy of Sciences 2005; 1041:248–255.

5. Gray CA, Bazer FW, Spencer TE. Effects of Neonatal Progestin Exposure on Female Reproductive Tract Structure and Function in the Adult Ewe1. Biology of Reproduction 2001; 64:797–804.

6. Bartol FF, Wiley AA, Spencer TE, Vallet JL. Early uterine development in pigs. Journal of Reproduction and fertility supplement 1993; 48:99–116.

7. Yan W, Chen J, Wiley AA, Crean-Harris BD, Bartol FF, Bagnell CA. Relaxin (RLX) and estrogen affect estrogen receptor α, vascular endothelial growth factor, and RLX receptor expression in the neonatal porcine uterus and cervix. Reproduction 2008; 135:705–712.

8. Yan W, Wiley AA, Bathgate RAD, Frankshun A-L, Lasano S, Crean BD, Steinetz BG, Bagnell CA, Bartol FF. Expression of LGR7 and LGR8 by Neonatal Porcine Uterine Tissues and Transmission of Milk-Borne Relaxin into the Neonatal Circulation by Suckling. Endocrinology 2006; 147:4303–4310.

9. Camp ME, Wiley AA, Boulos MB, Rahman KM, Bartol FF, Bagnell CA. Effects of age, nursing, and oral IGF1 supplementation on neonatal porcine cervical development. Reproduction 2014; 148:441–451.

10. Miller DJ, Wiley AA, Chen JC, Bagnell CA, Bartol FF. Nursing for 48 Hours from Birth Supports Porcine Uterine Gland Development and Endometrial Cell Compartment-Specific Gene Expression. Biology of Reproduction 2013; 88.

11. Bartol FF, Wiley AA, Miller DJ, Silva AJ, Roberts KE, Davolt MLP, Chen JC, Frankshun AL, Camp ME, Rahman KM, Vallet JL, Bagnell CA. Lactation Biology Symposium: lactocrine signaling and developmental programming. Journal of animal science 2013; 91:696.

12. Bartol FF, Wiley AA, Miller DJ, Silva AJ, Roberts KE, Davolt MLP, Chen JC, Frankshun A-L, Camp ME, Rahman KM, Vallet JL, Bagnell CA. LACTATION BIOLOGY SYMPOSIUM: Lactocrine signaling and developmental programming. Journal of Animal Science 2013; 91:696–705.

13. Chen JC, Frankshun A-L, Wiley AA, Miller DJ, Welch KA, Ho T-Y, Bartol FF, Bagnell CA. Milk-borne lactocrine-acting factors affect gene expression patterns in the developing neonatal porcine uterus. Reproduction 2011; 141:675–683.

14. Cai Y. Revisiting old vaginal topics: conversion of the Müllerian vagina and origin of the “sinus” vagina. International Journal of Developmental Biology 2009; 53:925–934.

15. Casey T, Harlow K, Ferreira CR, Sobreira TJP, Schinckel A, Stewart K. The potential of identifying replacement gilts by screening for lipid biomarkers in reproductive tract swabs taken at weaning. Journal of Applied Animal Research 2018; 46:667–676.

16. Casey T, Harlow K, Ferreira CR, Sobreira TJ, Schinckel A, Stewart K. The potential of identifying replacement gilts by screening for lipid biomarkers in reproductive tract swabs taken at weaning. Journal of Applied Animal Research 2018; 46:667–676.

17. Benevenga N, Steinman-Goldsworthy J, Crenshaw T, Odle J. Utilization of Medium-Chain Triglycerides by Neonatal Piglets: I. Effects on Milk Consumption and Body Fuel Utilization1. Journal of animal science 1989; 67:3331–3339.

18. Odle J, Benevenga N, Crenshaw T. Utilization of medium-chain triglycerides by neonatal piglets: II. Effects of even-and odd-chain triglyceride consumption over the first 2 days of life on blood metabolites and urinary nitrogen excretion. Journal of Animal Science 1989; 67:3340–3351.

19. Le Dividich J. Effect of fat content of colostrum on voluntary colostrum intake and fat utilization in newborn pigs. Journal Of Animal Science 1997; 75:707.

20. Su G, Zhou X, Wang Y, Chen D, Chen G, Li Y, He J. Effects of plant essential oil supplementation on growth performance, immune function and antioxidant activities in weaned pigs. Lipids Health Dis 2018; 17:139.

21. Wieland TM, Lin X, Odle J. Utilization of medium-chain triglycerides by neonatal pigs: effects of emulsification and dose delivered. Journal of animal science 1993; 71:1863–1868.

22. Bligh E, Dyer W. A rapid method of total lipid extraction and purification. Can J Biochem Physiol 1959; 37:911–917.

23. Chong J, Soufan O, Li C, Caraus I, Li S, Bourque G, Wishart DS, Xia J. MetaboAnalyst 4.0: towards more transparent and integrative metabolomics analysis. Nucleic acids research 2018.

24. Faraggi D, Reiser B, Schisterman EF. ROC curve analysis for biomarkers based on pooled assessments. Statistics in Medicine 2003; 22:2515–2527.

25. Xia J, Broadhurst D, Wilson M, Wishart D. Translational biomarker discovery in clinical metabolomics: an introductory tutorial. Metabolomics 2013; 9:280–299.

26. Forman BM, Tontonoz P, Chen J, Brun RP, Spiegelman BM, Evans RM. 15-Deoxy-Δ12,14-Prostaglandin J2 is a ligand for the adipocyte determination factor PPARγ. Cell 1995; 83:803–812.

27. Kliewer SA, Sundseth SS, Jones SA, Brown PJ, Wisely GB, Koble CS, Devchand P, Wahli W, Willson TM, Lenhard JM, Lehmann JM. Fatty acids and eicosanoids regulate gene expression through direct interactions with peroxisome proliferator-activated receptors α and γ. Proceedings of the National Academy of Sciences 1997; 94:4318–4323.

28. Delplanque B, Gibson R, Koletzko B, Lapillonne A, Strandvik B. Lipid Quality in Infant Nutrition: Current Knowledge and Future Opportunities. Journal of pediatric gastroenterology and nutrition 2015; 61:8–17.

29. Lanza I, Shoup DI, Saif LJ. Lactogenic immunity and milk antibody isotypes to transmissible gastroenteritis virus in sows exposed to porcine respiratory coronavirus during pregnancy. Am J Vet Res 1995; 56:739–748.

30. Drew MD, Bevandick I, Owen BD. Artificial rearing of colostrum-deprived piglets using iron chelators: The effects of oral administration of EDDHA with and without bovine or porcine immunoglobulins on piglet performance and iron-metabolism. Can. J. Anim. Sci. 1990; 70:655–666.

31. Bourne FJ, Newby TJ, Evans P, Morgan K. The immune requirements of the newborn pig and calf. Ann Rech Vet 1978; 9:239–244.

32. Davidson MH, Johnson J, Rooney MW, Kyle ML, Kling DF. A novel omega-3 free fatty acid formulation has dramatically improved bioavailability during a low-fat diet compared with omega-3-acid ethyl esters: the ECLIPSE (Epanova^®^ compared to Lovaza^®^ in a pharmacokinetic single-dose evaluation) study. Journal of clinical lipidology 2012; 6:573–584.

33. Hamosh M, Bitman J, Wood DL, Hamosh P, Mehta N. Lipids in milk and the first steps in their digestion. Pediatrics 1985; 75:146–150.

